# Multi-modal regulation of *C. elegans* hermaphrodite spermatogenesis by the GLD-1-FOG-2 complex

**DOI:** 10.1101/386250

**Authors:** Shuang Hu, Lauren E. Ryan, Ebru Kaymak, Lindsay Freeberg, Te-Wen Lo, Scott Kuersten, Sean P. Ryder, Eric S. Haag

## Abstract

Proper germ cell sex determination in *Caenorhabditis* nematodes requires a network of RNA-binding proteins (RBPs) and their target mRNAs. In some species, changes in this network enabled limited XX spermatogenesis, and thus self-fertility. In *C. elegans*, one of these selfing species, the global sex-determining gene *tra-2* is regulated in germ cells by a conserved RBP, GLD-1, via the 3’ untranslated region (3’UTR) of its transcript. A *C. elegans*-specific GLD-1 cofactor, FOG-2, is also required for hermaphrodite sperm fate, but how it modifies GLD-1 function is unknown. Germline feminization in *gld-1* and *fog-2* null mutants has been interpreted as due to cell-autonomous elevation of TRA-2 translation. Consistent with the proposed role of FOG-2 in translational control, the abundance of nearly all GLD-1 target mRNAs (including *tra-2)* is unchanged in *fog-2* mutants. Epitope tagging reveals abundant TRA-2 expression in somatic tissues, but an undetectably low level in wild-type germ cells. Loss of *gld-1* function elevates germline TRA-2 expression to detectable levels, but loss of *fog-2* function does not. A simple quantitative model of *tra-2* activity constrained by these results can successfully sort genotypes into normal or feminized groups. Surprisingly, *fog-2* and *gld-1* activity enable the sperm fate even when GLD-1 cannot bind to the *tra-2* 3’ UTR. This suggests the GLD-1-FOG-2 complex regulates uncharacterized sites within *tra-2*, or other mRNA targets. Finally, we quantify the RNA-binding capacities of dominant missense alleles of GLD-1 that act genetically as “hyper-repressors” of *tra-2* activity. These variants bind RNA more weakly *in vitro* than does wild-type GLD-1. These results indicate that *gld-1* and *fog-2* regulate germline sex via multiple interactions, and that our understanding of the control and evolution of germ cell sex determination in the *C. elegans* hermaphrodite is far from complete.

## Introduction

*C. elegans* hermaphrodites are essentially modified females, producing first sperm, then oocytes. These self-sperm allow uniparental reproduction, which has evolved at least three times in *Caenorhabdtis*. Selfing is likely to be adaptive in the patchy, ephemeral environments in which these animals reproduce (Kiontke et al., 2011). How did XX females evolve the ability to produce sperm? Decades of genetic and molecular analyses have identified a network of gene products required to promote XX spermatogenesis in *C. elegans*. Central to this network is *tra-2*, a global sex determiner that was first identified by Hodkgin and Brenner (1977) by its Transformer (Tra) null phenotype. In XX *tra-2* loss-of-function (*tra-2(lf)*) mutants, both the soma and germ cells of hermaphrodites adopt the male form. *tra-2* gain-of-function alleles also exist that have a feminization of germline (Fog) phenotype: XX animals make oocytes, but no sperm (Doniach, 1986; Goodwin et al., 1993). All such *tra-2(gf)* alleles map to a direct repeat element (DRE) in the 3’UTR of *tra-2*, and have little or no effect on male spermatogenesis. These data implicate the presence of a post-transcriptional negative regulator of *tra-2* in XX hermaphrodites that is essential for spermatogenesis (Puoti et al., 1997).

Like the *tra-2* DREs, *gld-1* is required for spermatogenesis in XX hermaphrodites, but not males, although its phenotypes are more pleiotropic (Francis et al., 1995a; Francis et al., 1995b). In addition to loss of XX sperm, *gld-1(null)* mutants have germ cell tumors of oocyte character, caused by meiotic failure and resumption of mitosis. GLD-1 is a STAR family RBP that is most abundant in early meiotic cells (Jones et al., 1996). It was identified in a molecular screen for its ability to bind the DREs (Jan et al., 1999), indicating it is the germline factor that binds the *tra-2* 3’UTR. Subsequent studies have confirmed the association of GLD-1 with the *tra-2* mRNA (Jungkamp et al., 2011; Wright et al., 2011) and revealed that GLD-1 likely binds its targets as a homodimer (Ryder et al., 2004).

A third component that is essential specifically for XX spermatogenesis is encoded by *fog-2*. Similar to *tra-2* gain-of-function mutants, *fog-2* null XX mutants are abnormal only in their lack of sperm, while spermatogenesis of XO male mutants is unaffected (Schedl and Kimble, 1988). This germline-feminizing effect requires *tra-2* function, suggesting *fog-2* acts upstream of *tra-2*. The *fog-2* gene was eventually cloned by virtue of a physical interaction between FOG-2 and GLD-1 (Clifford et al., 2000). FOG-2 possesses both an F-box and a novel GLD-1-binding domain, the latter distinguishing it from other members of the large F-box family of paralogs (Nayak et al., 2005). F-box proteins often link E3 ubiquitin ligases to their target proteins (Genschik et al., 2013; Ho et al., 2008). Shimada et al. (2006) found that a non-essential proteasome component (RPN-10) is required for XX sperm and *tra-2* repression body-wide. These results all are consistent with a model in which FOG-2 and GLD-1 evolved in the *C. elegans* lineage to cooperate to down-regulate *tra-2* activity (Figure 1), and that ubiquitination and/or the proteasome are somehow involved.

**Figure 1.**
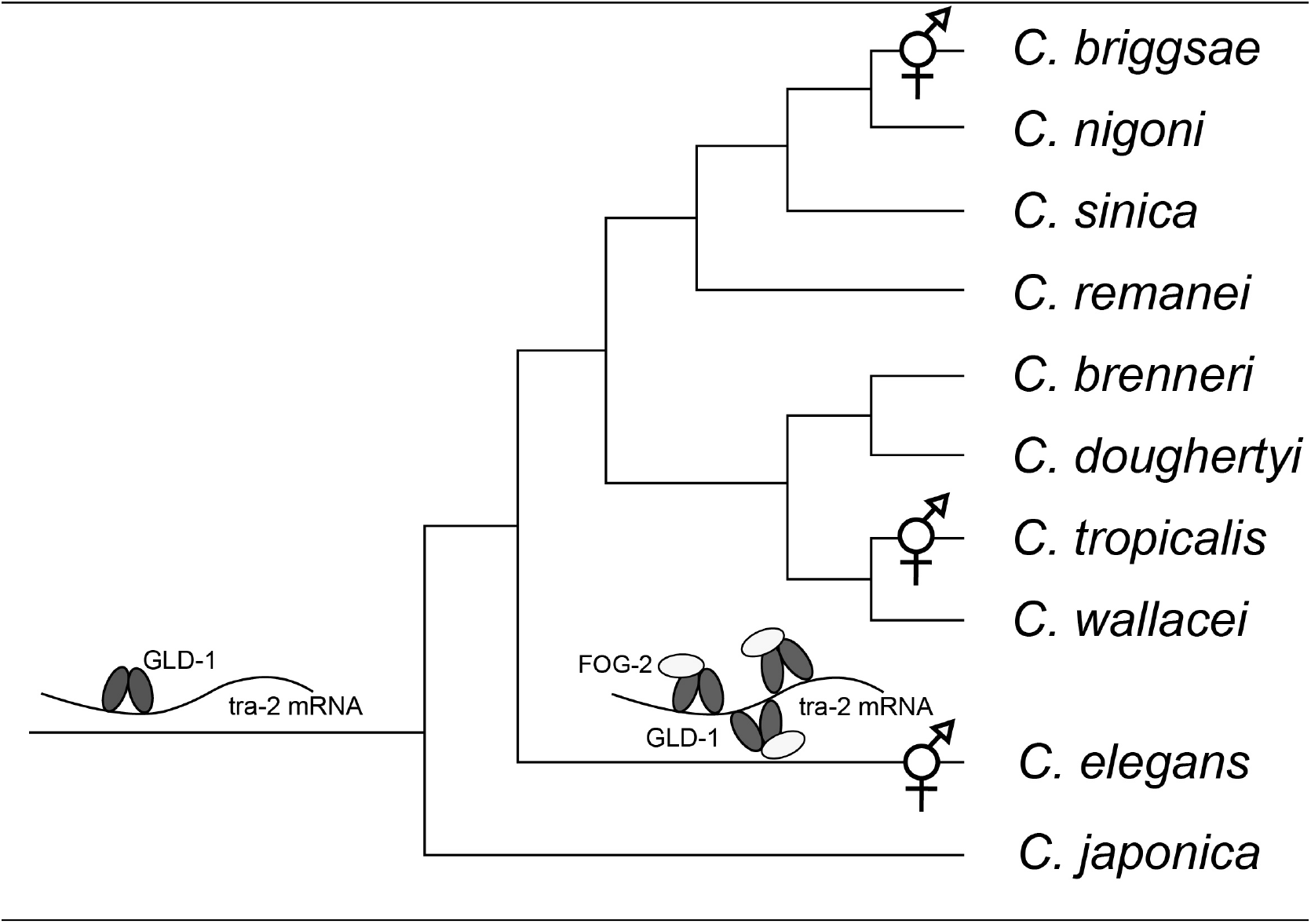
Evolutionary history of *gld-1* and *fog-2* in *Caenorhabditis* nematodes. A phylogeny (modified from Kiontke et al., 2011) showing the convergent evolution of selfing in the Elegans group species of *Caenorhabditis*. Derived aspects of *tra-2* regulation in *C. elegans* are depicted in cartoon form. The GLD-1-*tra-2* mRNA regulation was strengthened in XX hermaphrodites by 1) multimerization of GLD-1 binding elements in the *tra-2* mRNA’s 3’- untranslated region, or UTR (Beadell et al., 2011), and 2) recruiting the novel cofactor FOG-2 (Nayak et al., 2005). The convergently evolved hermaphrodite *C. briggsae* lacks these two features, and uses GLD-1 to opposite effect in germline sex determination (Beadell et al., 2011; Nayak et al., 2005).

More recent work has revealed insights into the mechanisms of GLD-1 function, with implications for its role in promoting hermaphrodite spermatogenesis. Ryder et al. (2004) first defined the consensus GLD-1 binding site in RNA targets. The extent of an mRNA’s association with GLD-1 is determined by the strength and number of these (and similar) sites (Galarneau and Richard, 2009; Wright et al., 2011), with most residing in 3’ UTR regions. The PAR-CLIP method, which identifies specific sites bound by RNA-binding proteins in native transcripts (Hafner et al., 2010), has been applied to *C. elegans* GLD-1 (Jungkamp et al., 2011). This revealed that GLD-1 binds near the start codon of some transcripts, suggesting that it may regulate interaction with the translation initiation machinery. While translation of GLD-1 target mRNAs is often repressed, this repression can also be accompanied by protection from nonsense mediated decay (Lee and Schedl, 2004) and stabilization (Scheckel et al., 2012). Thus, whether loss of *gld-1* function has a net positive or negative effect on target gene activity may vary from target to target.

GLD-1 protein is localized to the distal germ line of oogenic XX gonads, with greatest expression concentrated in the region marking entry into meiosis (Jones et al., 1996). GLD-1 expression is repressed in the distal germline stem cell niche by translational control by the FBF PUF family RBPs (Crittenden et al., 2002; Suh et al., 2009) and by *glp-1/Notch* signaling (Brenner and Schedl, 2016), such that cells upregulate GLD-1 as they leave the niche. The defects seen after loss of *gld-1* function (Beadell et al., 2011; Francis et al., 1995a; Francis et al., 1995b; Nayak et al., 2005) and the identity of mRNAs bound by GLD-1 (Beadell et al., 2011; Lee and Schedl, 2001; Ryder et al., 2004; Wright et al., 2011) in multiple *Caenorhabditis* species suggest that the ancestral role of *gld-1* was in the meiotic progression and differentiation of oocytes.

Comparative studies among *Caenorhabditis* species have revealed that sex determination mechanisms have rapidly evolved. *C. briggsae*, a convergently evolved hermaphroditic species, has a *tra-2* ortholog with a conserved female-promoting function (Kuwabara,1996; Kelleher et al., 2008). However, while *gld-1* is present in *C. briggsae*, its role in self fertility is distinct. RNAi knockdown of *gld-1* in *C. briggsae* results in a sperm-only (Mog) phenotype, unlike the feminization seen in *C. elegans* (Nayak et al., 2005). These RNAi results were confirmed with *Cbr-gld-1* loss-of-function mutants by Beadell et al. (2011). Despite these opposite loss-of-function sex phenotypes, a *Cbr-gld-1* transgene can fully rescue the *C. elegans gld-1* null mutant. This suggests a pivotal role for changes in target mRNAs in the evolution of GLD-1 function. Consistent with this, the *C. briggsae tra-2* 3’ UTR lacks the cluster of GLD-1-binding sites that are essential for proper *C. elegans tra-2* regulation (Beadell et al., 2011).

The *tra-2* ortholog of the male/female *C. remanei* has a conserved role in female sexual fate, and a GLD-1-like factor binds to the *C. remanei tra-2* 3’ UTR (Haag and Kimble, 2000). This suggests that key elements of the *C. elegans* system of germline sex regulation predate the evolution of selfing. However, despite large families of F-box proteins in all Caenorhabditis (Nayak et al., 2005; Thomas, 2006), only *C. elegans* has a *fog-2* paralog. Interestingly, *C. briggsae* evolved selfing independently (Kiontke et al., 2011; Kiontke et al., 2004), yet requires its own species-restricted F-box protein, SHE-1, for spermatogenesis in hermaphrodites (Guo et al., 2009).

A number of important questions remain unanswered about how XX spermatogenesis is regulated in *C. elegans*. Many of these relate to *tra-2, gld-1, fog-2* and the nature of their interactions. Does FOG-2 impact the steady-state transcript levels of GLD-1 targets, including *tra-2*? Can the inferred repression of germline TRA-2 levels by the combined action of GLD-1 and FOG-2 be directly visualized? Can the function of *fog-2* and *gld-1* be ascribed wholly to regulation of *tra-2* via its 3’ UTR? What impact do the missense changes in dominant Mog alleles of *gld-1*, which act like “super repressors,” have on GLD-1 function? Below we describe experiments that address these questions, some using tools not available when the genetic interactions between *tra-2, gld-1*, and *fog-2* were first characterized.

## Materials and Methods

### C. elegans strains

Strains used in this study:

N2 (reference wild-type)
JK574 *fog-2(q71)V*
JK992 *tra-2(e2020)II; unc-24(e138) fem-3(q95) dpy-20(e1282)IV*
JK3107 *fbf-1(ok91) fbf-2(704)/mIn1[mIs14 dpy-10(e128)]II*
TX189 *teIs1*[*oma-1p::oma-1::GFP* + *unc-119(+)*].
BS3156 *unc-13(e51) gld-1(q485)/hT2[dpy-18(h662)]I; +/hT2[bli-4(e937)]III*
CP139 and CP141 *nm75[tra-2::HA]II* (see below)

CP153 *nm75[tra-2::HA]II; fog-2(q71)V* was generated by crossing JK574 males with CP139 or CP141 hermaphrodites. Homozygosity of the HA-tagged endogenous *tra-2* allele was confirmed by PCR amplicon band shift; and of the *q71* point mutation by PCR and sequencing. In the presence of *nm75*, the Fog phenotype of *q71* is incompletely penetrant.

CP163 *unc-13(e51) gld-1(q485)/+I; nm75[tra-2::HA]II* was generated by crossing N2 males with CP141 hermaphrodites, then using F1 males to cross with phenotypically wild-type BS3156 hermaphrodites. Homozygosity of the HA-tagged *tra-2* was confirmed by PCR, and of *q485* by the gonad tumor and linked Unc phenotypes.

Triple mutant *tra-2(e2020) II; fem-3(q95) I; fog-2(q71) V* were generated by using *mIn1* as a marker on the second chromosome. Dpy *mIn1II; fog-2(q71) V* males were crossed with JK992 *tra-2(e2020) II; unc-24(e138) fem-3(q95) dpy-20(e1282) IV*, to yield wild-type like F1s. Non-green, Dpy Unc F2s F2 progeny are *tra-2(e2020); fem-3(q95)*, but of uncertain *fog-2* genotype. After phenotyping under a compound microscope, a subset were recovered and a PCR amplicon (produced with primers EH42 5’- CTTCACATGAAGCTTCAAAGC-3’ and EH43 5’- TTTGAGGGGAAAAATGTCCAATT-3’) was sequenced to genotype the point mutation of *fog-2(q71)*.

CP165 *unc-24(e138) fem-3(q95) dpy-20(e1282)IV; fog-2(q71) V* was created by crossing JK992 hermaphrodites with *fog-2(q71)* males, and singling many Dpy Unc F2 to create known *fem-3(q95)* founders. These F2 founders were allowed to lay eggs, and then PCR genotyped to eliminate families carrying *tra-2(e2020)*. Finally, *fog-2* genotypes of *fem-3(q95); tra-2(+/+)* families were determined by PCR and sequencing as described above. Because *fog-2(q71)* enhances the fertility of *fem-3(q95)* at permissive temperature (Schedl and Kimble, 1988), *q71* homozygotes were over-represented among healthy lines.

Most strains were kept at 20°C on NGM agar plates, except for those carrying *fem-3(q95)*, which is temperature-sensitive. Such strains were held at 15°C for propagation, and assayed for germline feminization by shifting embryos to 25°C and growing them to adulthood at this restrictive temperature.

### RNAseq comparisons

Nematodes were synchronized by allowing purified embryos to hatch in food-free M9 salts, and then moving arrested hatchlings to standard NGM plate media with *E. coli* OP50 bacterial food. Three independent samples of 100 N2 hermaphrodites and *fog-2(q71)* females were harvested on their first day of adulthood. To avoid potential changes in the oocyte transcriptome related to suppressed ovulation (Jud et al., 2008), only *fog-2* XX females that had successfully mated and were gravid were chosen. RNA was extracted in Trizol (Ambion) and purified as previously described (Thomas et al., 2012). cDNA was synthesized using either a 3’- anchored oligo-dT primer or random hexamers via the TotalScript kit (Illumina). As part of ongoing product development work, hexamer-primed libraries were generated using one of two alternative formulations (“standard” or “optimized”) of the reaction buffer for synthesis of the first cDNA strand. The combination of buffer and sequencing run that produced the greatest number of non-rRNA mapped reads was used for subsequent analyses, resulting in use of “optimized” libraries for all but replicate D of the *fog-2* samples. Libraries used unique barcodes to allow multiplexing.

Sequencing of the cDNA libraries was performed on an Illumina HiSeq instrument. 30-50 million pairs of 51-nucleotide reads were generated for each library. For the oligo-dT-primed libraries nearly all reads were derived from mRNAs (not shown). For hexamer-primed libraries, roughly 85% of reads were derived from rRNA. After de-multiplexing, reads were mapped to annotated genes of the N2 reference genome (release CE10) using TopHat (Trapnell et al.,2009). Expression levels and statistical tests for significant differences between genotypes was assesses using the program *cuffdiff*, within the Cufflinks package (Trapnell et al., 2010). Results reported here were obtained using the per-condition dispersion option. Expression measurements, statistical tests, and software details for this study are available from the National Center for Bioinformatics (NCBI)’s Gene Expression Omnibus via accession number GSE110998.

### CRISPR genomic modification

Hemagglutinin (HA) epitope tag was knocked into the endogenous *tra-2* locus by CRISPR-Cas9 genomic editing. The 9 codons encoding the HA tag was inserted so as to add epitope near the C-terminus of TRA-2, 4 codons before the stop codon. A single guide RNA with the targeting sequence GAAGGCGACCTATCAGACCCAG was *in vitro* transcribed using the SP6 Megascript system (Invitrogen-Ambion). Oligonucleotide-based transcription templates were designed and synthesized according to the method of Gagnon et al. (2014), with the exception of using Taq DNA polymerase for the annealed oligonucleotide fill-in reactions. The repair oligo sequence (Ultramer, IDT) used for homology-directed repair was 5’ gatgaagcacgggaaggcgacctatcagacT ACCCAT ACGA

CGTTCCAGACTACGCCccagaggtttaaaatgtctgtttcctttttcag 3’, where uppercase sequences indicate the HA-encoding insertion, and underlined nucleotides indicate the Cas9 guide RNA sequence.

Well-fed young adult worms were injected with the mixture of 800 ng/μl nuclear localization sequence-conjugated Cas9 protein (PNABio), 900 ng/μl sgRNA, and 60nM repair oligo. Edited heterozygous F1 were identified by single worm PCR among the offspring within 8-24 hours time window after injection, using primers flanking the insertion region, looking for a PCR product shift from 203nt to 230nt. *gld-1 RNA interference*

An in vitro transcription template for production of *gld-1* dsRNA was synthesized by PCR using the forward primer 5’-

TAATACGACTCACTATAGGGTGGATGCAAGATTATGGTCCGAGG-3’ and reverse primer 5’-TAATACGACTCACTATAGGGGACCAGGGAATGAGTTGACAAACG-3’ (T7 promoter sequence underlined). The PCR product was purified with a QIAquick PCR purification kit (Qiagen) and 800 ng was used for *in vitro* transcription using a Megascript T7 kit (Ambion) following the factory protocol. Young adult JK992 and N2 hermaphrodites were injected in the germline or gut at a concentration of 2000 ng/μl, and allowed to recover for 6 hours at 15°C. They were then moved to new plates and allowed to lay for 24 hours, after which embryos were shifted to 25°C or left at 15°C as controls.

### Immunoblots and Immunohistochemistry

For immunoblots, 200 worms of each strain were boiled in 2x Laemmli Loading buffer (Bio-Rad), and the entire lysate was fractionated on a 12% SDS-polyacrylamide gel and transferred to nitrocellulose membranes using standard conditions (Harlow and Lane, 1999). Immunoblots used the rat anti-HA monoclonal antibody 3F10 (Roche) 1:100 as the 1^st^ antibody, followed by goat anti-rat IgG-HRP sc-2006 (Santa Cruz) 1:500 as the 2^nd^ antibody. Actin was detected by anti-actin antibody C4 1:1000 (sc-47778), followed by bovine anti-mouse HRP sc-2375 (Santa Cruz) 1:5000. HRP activity was detected on blots using chemiluminescence (SuperSignal West Pico substrate, Thermo Scientific).

For immunohistochemistry, animals were dissected in 0.25 levamisole in PBS, either in 4-well polycarbonate tissue culture trays (for L4 larvae and adults) or on poly-L-lysine-subbed microscope slides (for L3 larvae). Dissected worms were fixed in 3% formaldehyde in 0.1M K_2_HPO_4_, followed by dry methanol. Samples were rehydrated and stained with rat anti-HA 3F10 (Roche) 1:100, followed by Alexa Fluor goat anti-rat 555 (Molecular Probes Invitrogen) 1:100, both in PBS with 0.1% Tween. Nuclei were stained with 100 μg/ml Hoechst 33258 just prior to imaging. Images for L4 and adult samples were captured with Zen software on a Zeiss LSM 710 confocal microscope, at 6% laser power and default (1.0) gain, using 10x or 40x oil objectives. Images for L3 larvae were captured with a Leica SP5X confocal microscope, using a 40x oil objective with 3x digital zoom. The 405 nm laser was at 15% power with gain of 648. 555 nm excitation was achieved with a white laser source at 70% power, with the 554nm line at 60% power and 943 gain.

To quantify and compare germline TRA-2:HA expression, a region of at least 2000 pixels from a confocal section that included the distal tip cell was quantified using the histogram tools of Zen (for L4 and adult samples) or ImageJ (for L3 samples) image analysis packages. Samples compared were dissected and stained in parallel using identical reagents on the same day, and imaged in the same session using identical laser and image capture settings.

### GLD-1-RNA binding assays

GLD-1 expression plasmids: The G248R and G250R mutant derivatives of pHMTc-GLD(135- 336) (Ryder et al., 2004) were produced by QuikChange mutagenesis (Agilent Genomics, Santa Clara, CA) using primers 5’-CCAGA TTATG GTCCG AAGAA AGGGA TCAAT GC-3’ and 5’-GCATT GATCC CTTTC TTCGG ACCAT AATCT TG-3’ or primers 5’-GGTCC GAGGA AAGAG ATCAA TGCG GG-3’ and 5’-CCCGC ATTGA TCTCT TTCCT CGGAC C-3’, respectively. Sequencing was used to validate the presence of the desired mutations.

Protein purification: The plasmids encoding pHMTc-GLD(135-336) or mutations thereof were transformed into *Escherichia coli* strain BL21(DE3). All three maltose-binding protein-GLD-1 fusions were purified as previously described (Ryder et al., 2004). Briefly, bacterial cultures were induced at mid log phase for three hours using 1 mM isopropyl 1-thio-β-D-galactopyranoside (IPTG). After harvesting, the cell pellets were lysed, then soluble recombinant protein was purified across an amylose affinity column (New England Biolabs) followed by HiTrap Q and HiTrap S ion exchange columns (GE Healthcare, Wauwatosa, WI) at 4°C. Relative purity of the protein was assessed by SDS-PAGE, and the concentration estimated from the absorption of 280 nm UV light using extinction coefficients calculated from the sequence composition with Expasy Protparam (Wilkins et al., 1999). Purified proteins were dialyzed into storage buffer (50 mM Tris pH 8.0, 20 mM NaCl, 2 mM DTT) and kept at 4°C until use.

Binding assays: Electrophoretic mobility shift assays using purified recombinant MBP-tagged GLD-1 and mutant variants were performed essentially as described (Farley and Ryder, 2012). The target RNA sequence (RNA: 5’-UUUUU CUUAU UCUAG ACUAA UAUUG UAAGC U-3’-Fl), comprising a fragment of the *glp-1* 3’UTR previously shown to bind to GLD-1 (Farley and Ryder, 2012), was chemically synthesized and labeled at the 3’end with fluorescein amidite (FAM) by Integrated DNA Technologies (IDT, Coralville, IA). Varying concentrations of purified GLD-1, GLD-1 (G248R) and GLD-1 (G250R) were equilibrated with 3 nM of labeled RNA in equilibration buffer (0.01% IGEPAL, 0.01 mg/ml tRNA, 50 mM Tris, pH 8.0, 100 mM NaCl, 2mM DTT) for 3 hours. After equilibration, bromocresol loading dye was added, the reaction mixture was loaded onto a 5% native polyacrylamide gel in 1X TBE, and allowed to run for 2 hours at 120 volts. The gels were quantified by determining the pixel intensity of the RNA bound species and the total of all RNA species to give the fraction bound of RNA relative to the background using Image Gauge (Fujifilm, Tokyo, Japan). The graph of fraction bound against the protein concentration was then fit to the Hill equation to determine the apparent binding affinity and the Hill coefficient.

## Results

### Impact of fog-2 on the transcriptome

If GLD-1 regulates protein expression at the translational level, and FOG-2 functions via its association with GLD-1, then loss of FOG-2 may not affect steady-state levels of mRNAs targeted by GLD-1. Alternatively, the demonstrated impact of *gld-1* on target stability noted above would predict that some targets may show altered transcript levels. We therefore used RNAseq to compare the abundance of mRNAs extracted from gravid young adult N2 and *fog-2(q71)* XX worms, which are genetic nulls for *fog-2* function (Schedl and Kimble, 1988). Because GLD-1 binding sites can regulate the extent of *tra-2* mRNA polyadenylation (Thompson et al., 2000), and because deadenylation can enhance mRNA turnover (Alonso, 2012), separate sequencing libraries produced from cDNA synthesized with oligo-dT and random hexamer primers were generated for each sample.

In the oligo-dT-primed experiment, we found 235 of 19,291 transcripts (1.2%) were significantly differentially expressed (FDR = 0.05) between the wild-type and *fog-2(q71)* samples (**Figure 2, Suppl. Table 1**). 14 of these encode paralogous major sperm proteins (MSPs), sperm-specific cytoskeletal and signaling proteins (Burke and Ward, 1983; Ward et al., 1988). All 14 showed roughly four-fold reduced expression in *fog-2(q71)*, as did the sperm-specific family S gene *sss-1* and the msp-related *msd-4*. The differentially expressed *msp* genes represent half of the total (Smith, 2014), far more than the one case expected by chance. Absent from the set of differentially expressed transcripts was *tra-2* (mean FPKM values in wild-type and *fog-2(q71)* of 48.9 and 49.6, respectively). Furthermore, no known regulators of sex-determination were differentially expressed, with the exception of *fog-2* itself. The four-fold reduction of *fog-2* mRNA abundance in *fog-2(q71)* homozygotes indicates that the *q71* mutation, a premature stop (Clifford et al. 2000), may induce nonsense-mediated decay.

**Figure 2.**
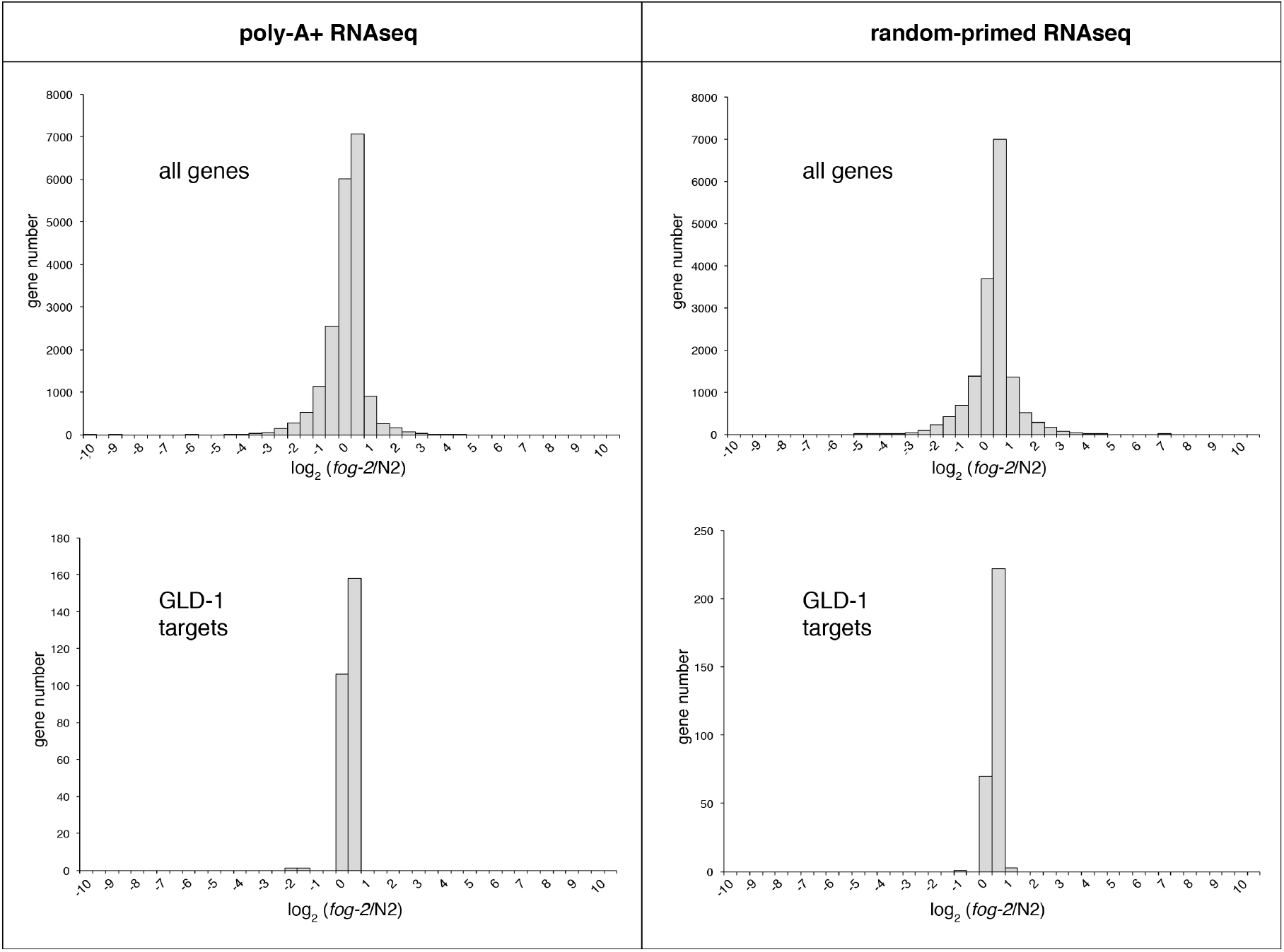
Steady-state mRNA levels in wild-type vs *fog-2(q71)* mutants. Top panels: Distributions of the fog-2/wild-type (N2) expression ratios for all transcripts consistently detected by RNAseq in young adult XX transcriptomes. Sequencing libraries were primed with oligo-dT (left) or random hexamers (right). Lower panels: Distributions of the fog-2/wild-type (N2) expression ratios for the 302 GLD-1 targets identified by both Wright et al. (2011) and Jungkamp et al. (2011). Data summarized from Supplemental Dataset 1.

**Table 1.** Quantitative model of germline *tra-2* activity. *T* represents the activity of one wild-type *tra-2* locus without trans-acting repression. *H, F*, and G represent factors (ranging from 0-1) that adjust *T* downward in response to the HA epitope tag of *tra-2(nm75)*, activity of *fog-2(+)*, or activity of *gld-1(+)*, respectively. *E* represents a factor (>1) by which *T* is increased in the *tra-2(e2020)* dominant Fog allele. Because reduction of *gld-1* and *fog-2* function impact sexual fate independently and in *e2020* mutants (see Tables 2 and 3), *F* and *G* are retained as factors even when combined with *tra-2(e2020)* alleles.

| Genotype | Germline phenotype | *tra-2* activity | value for $F=G=H=0.5$<br>$E>2$ |
| --- | --- | --- | --- |
| Wild-type | Hermaphrodite | $2TFG$ | $T/2$ |
| *tra-2(null)/+* | Hermaphrodite<sup>a</sup> | $TFG$ | $T/4$ |
| *tra-2:HA* | Hermaphrodite<sup>b</sup> | $2TFGH$ | $T/4$ |
| *tra-2(null)/+; fog-2(null)* | Hermaphrodite<sup>c</sup> | $TG$ | $T/2$ |
| *tra-2:HA; fog-2(null)* | Hermaphrodite<sup>b</sup> | $2TGH$ | $T/2$ |
| *tra-2:HA/+; fog-2(null)* | Fog<sup>b</sup> | $TG+TGH$ | $3T/4$ |
| *fog-2(null)* | Fog<sup>c</sup> | $2TG$ | $T$ |
| *gld-1(null)* | Fog (Tum)<sup>d</sup> | $2TF$ | $T$ |
| *tra-2(e2020)/+* | Fog<sup>e</sup> | $TEFG+TFG$ | $>3T/4$ |
| *tra-2(e2020)/tra-2(null)* | Fog<sup>e</sup> | $TEFG$ | $>T/2$ |
| *tra-2(null)* | Mog<sup>a</sup> | 0 | 0 |

<sup>a</sup> Hodgkin and Brenner (1977) <sup>b</sup> This study <sup>c</sup> Schedl and Kimble (1988) <sup>d</sup> Francis et al. (1995a) <sup>e</sup> Doniach (1986)

**Table 2.**
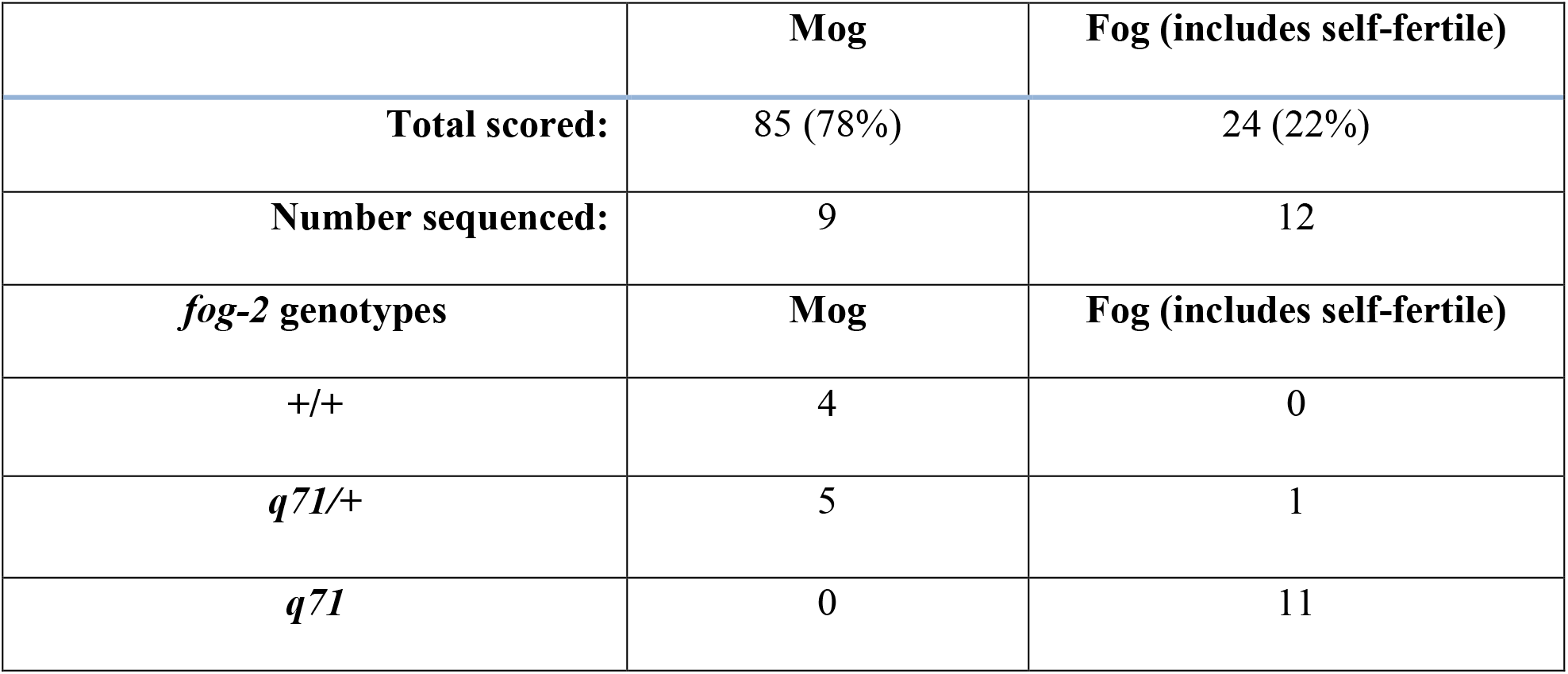
Evidence for a *tra-2* 3’ UTR-independent role for *fog-2*. Families of *tra-2 (e2020); fem-3 (q95)* mutants segregating for *fog-2(q71)* were grown at 25° C and scored for the presence of oocytes with a compound microscope (“total scored”). A subset of animals thus scored were recovered and genotyped at *fog-2* by single-worm PCR and sequencing.

|  | **Mog** | **Fog (includes self-fertile)** |
| --- | --- | --- |
| **Total scored:** | 85 (78%) | 24 (22%) |
| **Number sequenced:** | 9 | 12 |
| ***fog-2* genotypes** | **Mog** | **Fog (includes self-fertile)** |
| +/+ | 4 | 0 |
| *q71*/+ | 5 | 1 |
| *q71* | 0 | 11 |

**Table 3.**
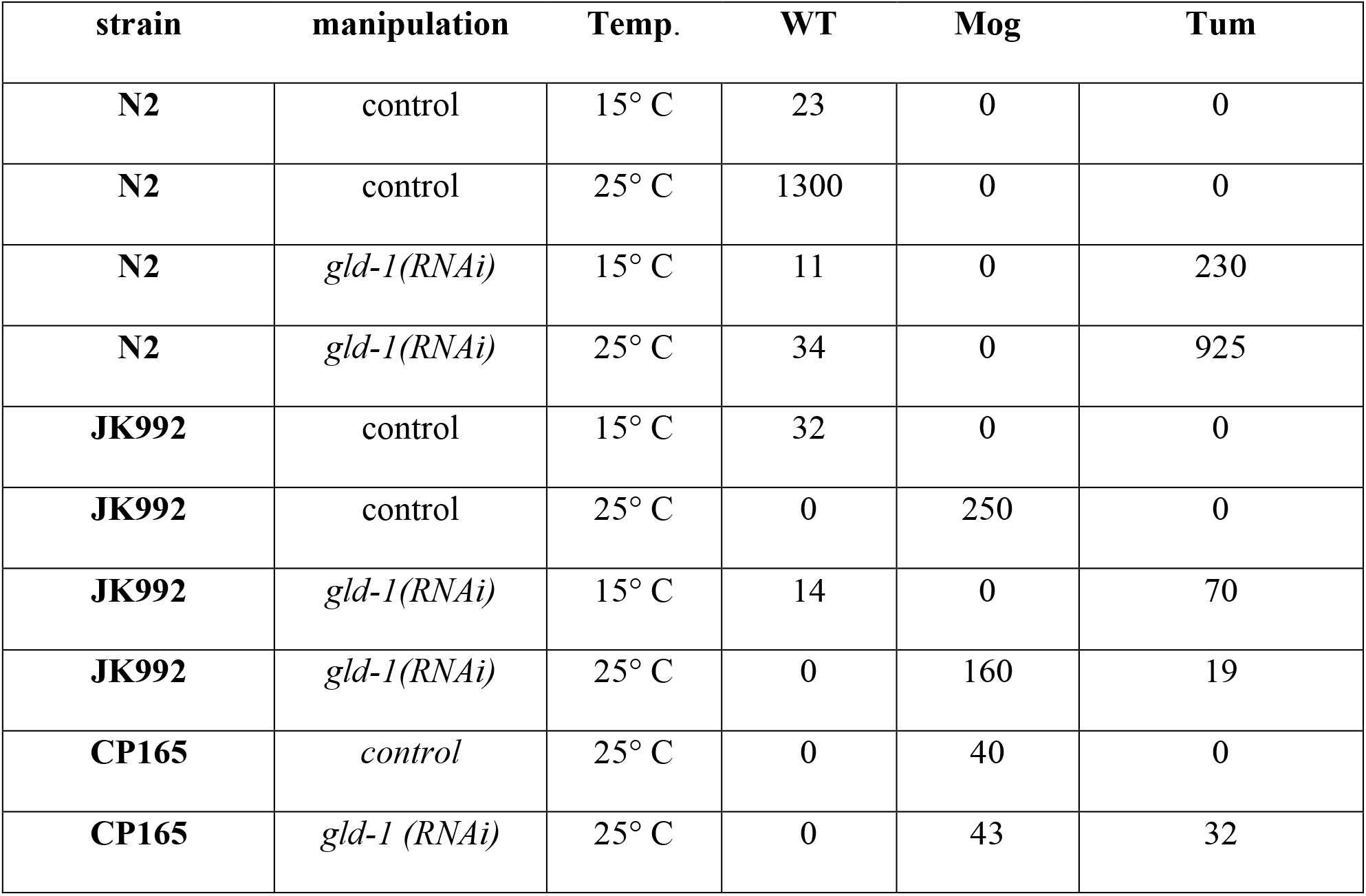
Genetic interactions between *gld-1(lf)* and *fog-2(lf), tra-2 (gf)*, and *fem-3 (gf)* Progeny of control and *gld-1(RNAi)* hermaphrodite injectees from the indicated strains were raised at either 15° or 25° C until young adulthood, and then scored with DIC microscopy. JK992 *is tra-2(e2020gf), fem-3(q95gf)*, and CP165 is *fem-3(q95); fog-2(q71)*, both in the N2 background.

In the RNAseq libraries produced using random hexamer primers (**Figure 2, Supplemental Table 1**), the burden of rRNA fragments reduced the number of genes whose expression could be consistently quantified to less than 10,000. Using the same significance threshold of FDR=0.05, 81 transcripts were judged as differentially expressed (0.08%). Reminiscent of the dT-primed data, the most significantly differentially expressed transcript was *fog-2*. In contrast with the oligo-dT-primed data, however, no *msp* paralogs were judged to significantly differentially expressed. However, for all 14 of the *msp* genes whose expression was detected in both genotypes in the random-primed libraries, mean expression was lower in the *fog-2* samples. This is highly unlikely to occur by chance given the overall symmetrical distribution of *fog-2*/N2 expression ratios (sign test P=.00006), and is therefore not a clear contradiction of the result from the dT-primed libraries.

Because FOG-2 lacks an RNA-binding domain, any direct role it may have in *tra-2* regulation likely relies on its association with GLD-1 (Clifford et al., 2000). We therefore examined the abundance of a set of 304 GLD-1-associated transcripts that were identified in two independent studies (Jungkamp et al., 2011; Wright et al., 2011). In both dT-primed and random-primed datasets, GLD-1 targets appear typical of the transcriptome as a whole, and are not enriched in differentially expressed transcripts (**Figure 2**). For the dT-primed case, two GLD-1 targets were judged to be differentially expressed (FDR=0.05), *gipc-1* and *R09E10.6*. This small number is similar to the number expected by chance (304 GLD-1 targets x 1.2% overall differential expression = 3.6). *gipc-1* encodes a PDZ domain-containing protein implicated in clearing of protein aggregates (Hamamichi et al., 2008), but with no known germ cell function. *R09E10.6* is a nematode-specific protein enriched in germ cells that may function in mitochondria (Sakamoto et al., 2012). In the random-primed dataset, none of the GLD-1 targets were judged to be differentially expressed. Overall, it appears that loss of *fog-2* has only subtle impacts on the transcriptome that are not biased towards GLD-1 targets or sex-determination genes. The similar results of the two libraries also indicate that FOG-2 does not regulate GLD-1 target polyadenylation.

### Expression of TRA-2 in wild-type and mutant contexts

To examine the regulation of TRA-2, we created an epitope-tagged allele. The two predicted TRA-2 isoforms are a full-length (~170 kDa) membrane protein (TRA-2A), transcribed from a 4.7kb mRNA, and a shorter TRA-2B protein (**Fig. 3a**). TRA-2B is translated directly in germ cells from a 1.8kb mRNA (Kuwabara et al., 1998), resulting in an ~55 kDa cellular protein. TRA-2B also corresponds to the intracellular domain of TRA-2A (i.e. “TRA-2ic”) after TRA-3 cleavage (Sokol and Kuwabara, 2000). To detect both TRA-2 isoforms, we added the small (9 amino acid) hemagluttinin (HA) epitope tag to endogenous TRA-2 using CRISPR genomic editing, inserting HA four amino acids before the stop codon. The resulting *tra-2:HA(nm75)* hermaphrodites are self-fertile and appear normal. TRA-2:HA is detected in larval and adult hermaphrodites (**Fig. 3b**), but not in males, consistent with their greatly reduced transcript levels (Okkema and Kimble, 1991). In anti-HA immunoblots, only the 55 kDa TRA-2B/ic isoform was detected.

**Figure 3.**
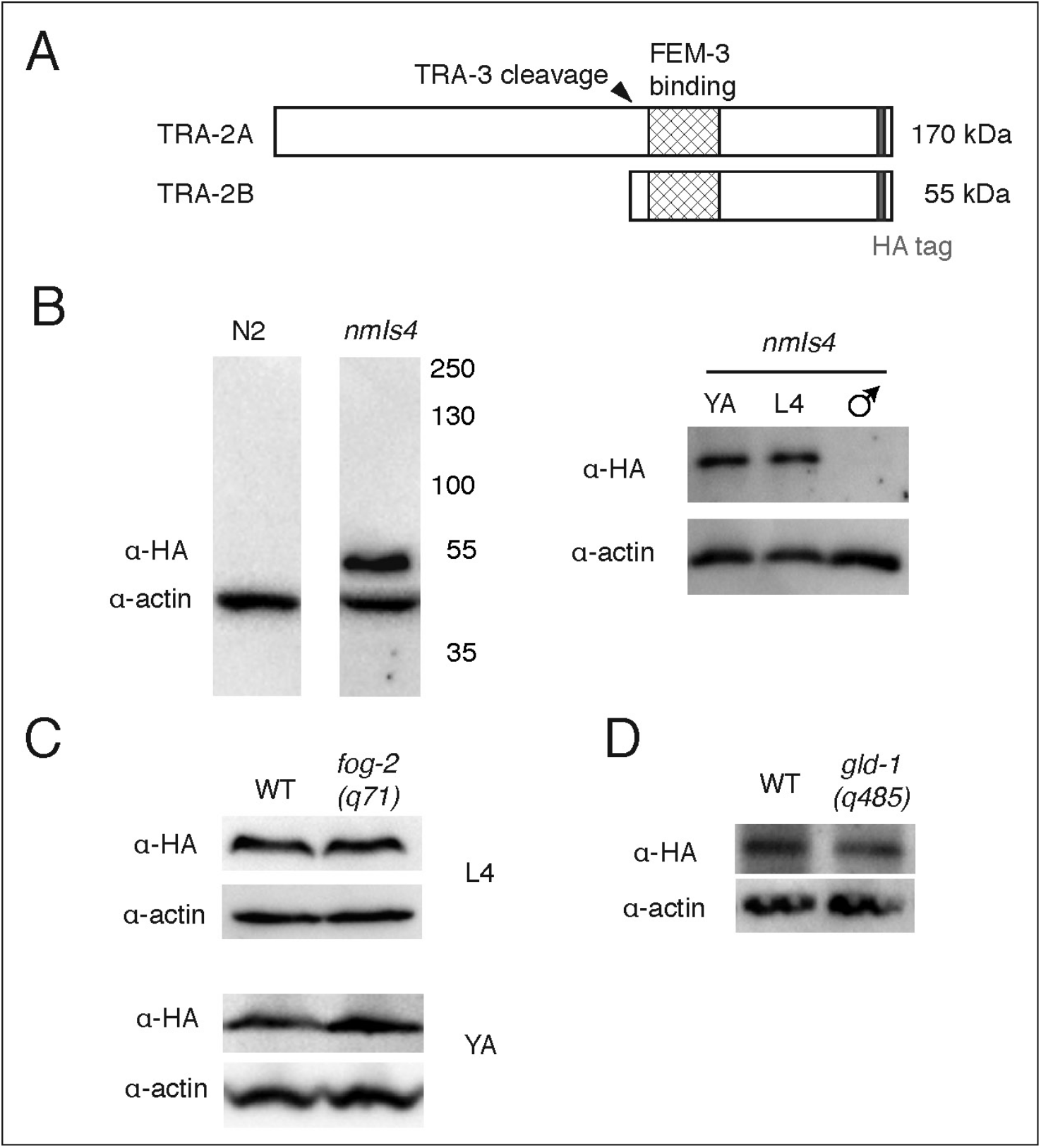
Impact of *tra-2* repressors on overall TRA-2B expression. A. Schematic of the two TRA-2 protein products, TRA-2A (a membrane protein) and TRA-2B (which lacks transmembrane domains). The HA epitope marks both forms near the carboxy terminus. B. Anti-HA antibodies detect TRA-2B:HA in lysates from XX *tra-2(nm75)* animals, but not in control N2 XX (left) nor XO *nm75* males (right). The full-length membrane protein TRA-2A:HA was not detected in either sex. C. TRA-2B:HA expression compared between wild-type XX hermaphrodites and *fog-2(q71)* XX mutant females at L4 (top) and young adult (bottom) stages.D. Comparison of TRA-2B:HA expression in lysates from adult XX wild-type (WT) and *gld-1(q485)* null mutants.

To reveal impacts of germline repressors of *tra-2* on TRA-2:HA expression, we first crossed the *tra-2:HA(nm75)* allele into the null *fog-2(q71)* allele (Schedl and Kimble, 1988). Hermaphrodite *fog-2(q71)* mutants do not produce sperm (the Fog, or feminization of germ line, phenotype), while males are unaffected. Surprisingly, 35% (N=40) of *tra-2:HA(nm75); fog-2(q71)* double mutants were self-fertile, and the strain can be propagated without any males present. Thus, the HA-tagged TRA-2 is a partial suppressor of *fog-2(q71)*, indicating that insertion of even the small (9-residue) HA tag encoded by *nm75* produces a mild reduction-of-function variant.

Because *fog-2* and *gld-1* are implicated in the suppression of *tra-2* translation, we hypothesized that TRA-2:HA expression would be increased in both *fog-2* and *gld-1* null mutant backgrounds. However, anti-HA immunoblots (**Fig. 3c, d**) revealed no elevation of TRA-2:HA signal in animals homozygous for null mutations in either *fog-2 (q71)* or *gld-1(q485)*. Thus, at the whole-worm level, TRA-2 expression was not detectably increased when *fog-2* or *gld-1* activities are absent.

### A quantitative model of tra-2 regulation in the hermaphrodite germ line

TRA-2 is a dose-sensitive protein (Kuwabara and Kimble, 1995), and both gain-of-function and loss-of-function *tra-2* alleles have defects in reproduction (Doniach, 1986; Hodgkin and Brenner, 1977). The precise level of *tra-2* activity thus appears to be crucial for germline sex determination (Puoti et al., 1997). In keeping with this, the *tra-2(nm75)* HA-tagged allele suppresses the *fog-2(q71)* Fog phenotype, yet does not reduce *tra-2* function enough to overtly masculinize an otherwise wild-type hermaphrodite. Similarly, one copy of the *tra-2(e1095)* strong loss-of-function allele also suppresses *fog-2(q71)* (Schedl and Kimble, 1988). *tra-2* null mutations are completely recessive (Hodgkin and Brenner, 1977), indicating that 50% of usual activity is adequate for development of a hermaphrodite germline. These observations suggest there is a range of *tra-2* activities that can allow the sperm-then-oocyte pattern. The multiple factors involved justify formalization of a quantitative model.

If the haploid dose of unmodified germline *tra-2* activity is defined as T, then the operational *tra-2* activity could be determined by functional copy number (0, 1, or 2) reduced by repressive factors (i.e. between 0 and 1), imparted by *fog-2 (F), gld-1 (G)*, or the HA epitope tag (H). Similarly, the *tra-2* 3’ UTR deletion allele, *e2020* (Doniach, 1986; Goodwin et al., 1993), augments *T* by a factor *E* (where E>1). **Table 1** presents expressions for various situations using these terms, assuming that a single copy of a negative regulator is sufficient for full function (i.e. they are dominant), that the activity of the two *tra-2* alleles is regulated independently and additively, and that *E, F, G*, and *H* are independent of each other. Justification for this last assumption comes from data presented in Tables 2 and 3, but is at best an approximation.

Despite its limitations, the above model supports some inferences. For example, the inferred *tra-2* activity in the wild-type case, *2TFG*, yields a hermaphrodite, while the *tra-2(e2020); tra-2(null)* expression, *TEFG*, is Fog. Since *tra-2* activity must be greater in the latter, it implies that E>2, or that the effect of the *e2020* 3’ UTR deletion is to at least double the basal level of *tra-2* activity. Similarly, though the repressive parameters *F, G*, and *H* are unknown, evaluating each the expression of *tra-2* activity for each genotype in Table 1 with *E>2* and *F=G=H=1/2* (right column) produces a clean separation between hermaphroditic and Fog genotypes, with hermaphrodite activity of up to *T/2* (for wild-type) and Fog activity as low as 3T/4 (for *tra-2:HA/+; fog-2(null))*. This discrimination between the two phenotypes is seen for any value between 1/3 and 1 when *F = G = H*. Allowing the values of *F, G*, or *H* to diverge can either increase or decrease the extent to which estimates of *tra-2* activity in the two phenotypic classes overlap. Given the data presented in Figure 6 below (which indicate that *gld-1* repression is greater than that of *fog-2)*, the effect of lowering *G* below *F* and *H* is worth noting (**Fig. 4**). Setting G=0.1, complete separation is seen when *F* and *H* are 0.5 or greater. The model is therefore robust to observed differences in repressor activity. More generally, it also highlights the bistable nature of germline sex determination, in which a continuous range of activity states produces only two discreet phenotypes.

**Figure 4.**
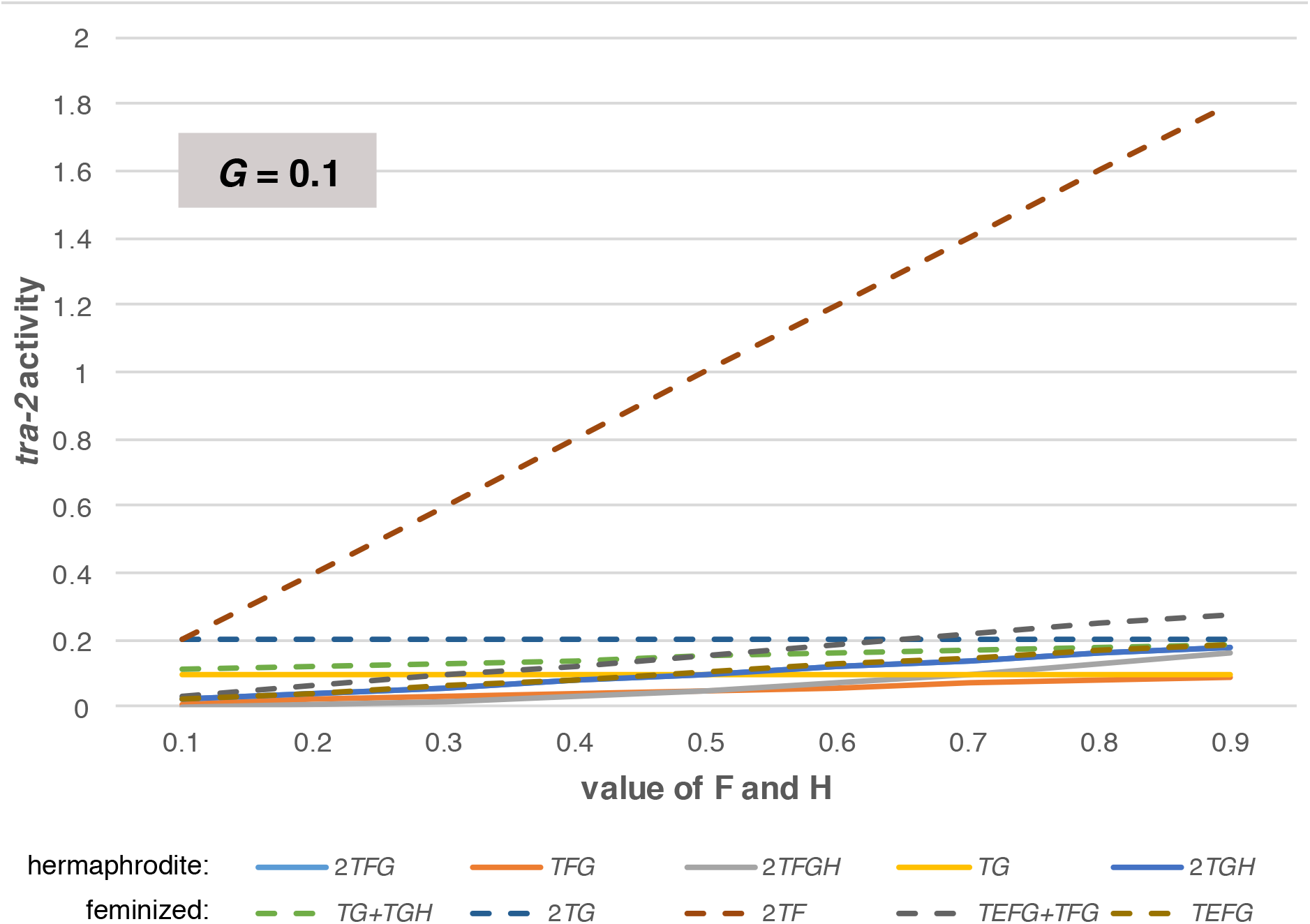
Modeling *tra-2* activity with strong *gld-1* repression. Each line corresponds to a hermaphroditic (solid) or feminized (dashed) genotype in Table 1. In calculating the expression for *tra-2* activity, the strength of *tra-2(e2020)* activation, E, is fixed at 2.1, and the parameter for *gld-1* repression, *G*, is fixed at 0.1 (i.e. very strong repression). The parameters for repression by *fog-2* and the *tra-2(nm75)* HA tag (*F* and H, respectively) are equal but allowed to vary along the X-axis as indicated. Note the separation of phenotypic classes by inferred *tra-2* activity when *F* = *H* > 0.5.

### TRA-2B:HA is detected in somatic nuclei, but not germ cells

The TRA-2:HA tagged allele described above *(nm75)* was next used to determine in which tissues TRA-2B is expressed, with particular emphasis on gonads. TRA-2B:HA was detected exclusively in the nuclei of dissected tissues of larvae and adults, including those of the intestine and somatic gonad (sheath and distal tip cells (**Fig. 5**). Nuclear localization of TRA-2B in somatic cells is consistent with previous studies using antibodies (Shimada et al., 2006) and reporter transgenes (Lum et al., 2000; Mapes et al., 2010). As judged by northern blotting (Kuwabara et al., 1998) and *in situ* hybridization (Kohara, 2005), *tra-2* transcripts are especially abundant in the germ line of adult hermaphrodites. Surprisingly, however, germ cell expression of TRA-2:HA was not consistently above background levels. Similar lack of germline expression was also reported with a polyclonal antibody raised against TRA-2B (Shimada et al., 2006). The TRA-2B expression observed in total worm lysates (**Fig. 3**) is therefore contributed almost completely from somatic tissues.

**Figure 5.**
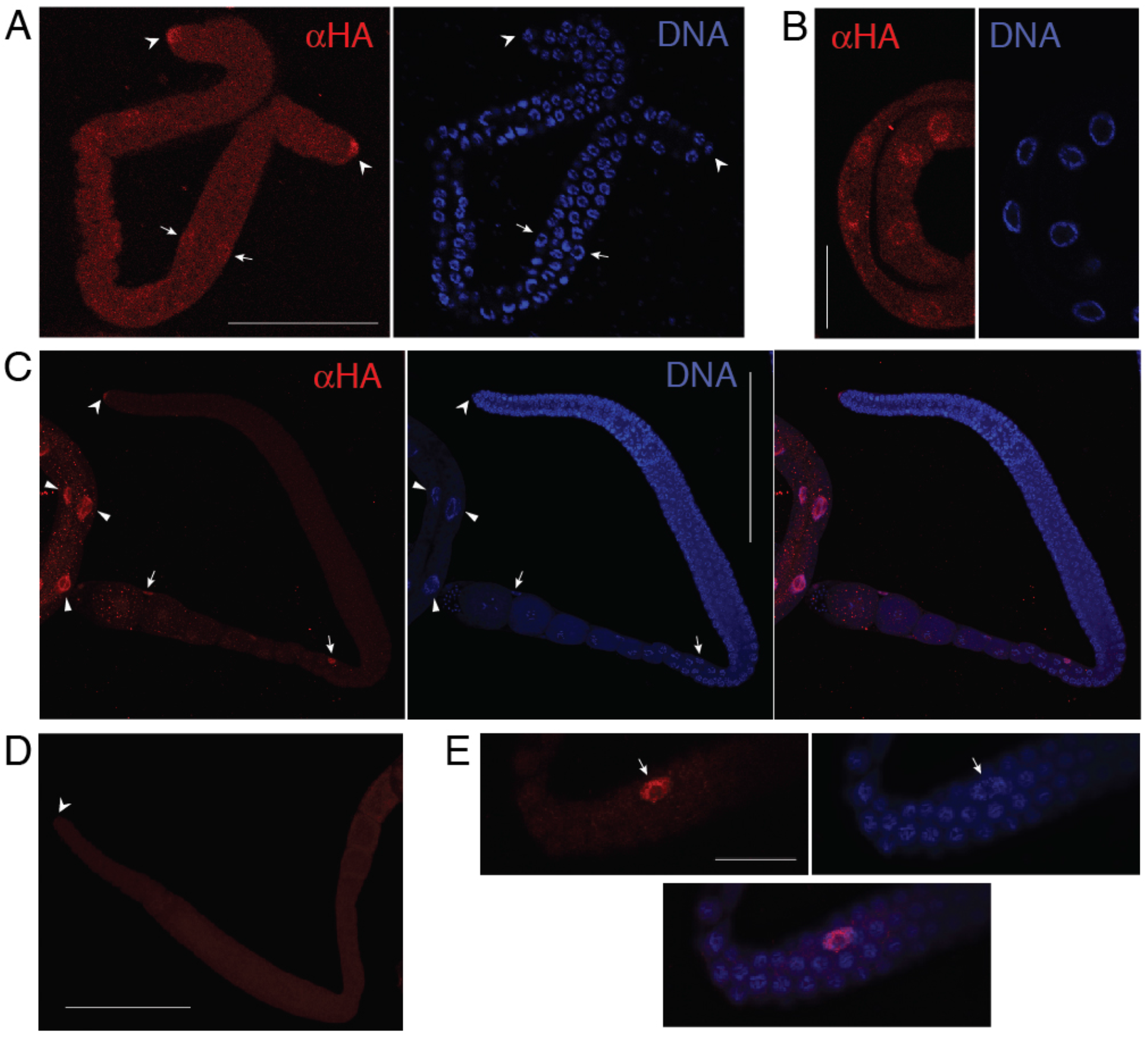
TRA-2:HA expression in wild-type animals. A, B. Confocal micrographs of dissected L3 larval hermaphrodite gonad (A) and intestine (B) showing TRA-2:HA localization (a-HA, left), Hoechst 33258 dye (DNA, right). TRA-2B:HA is detected in the nuclei of the distal tip cells (arrow heads) and more weakly in more proximal somatic cells (arrows). Scale bar is μm in A, 25 μm in B. C. Confocal micrograph of dissected adult hermaphrodite, showing TRA-2:HA localization (a-HA, left), Hoechst 33258 dye (DNA, center), and the two channels merged (right). TRA-2B:HA is detected in the somatic gonad, including the nuclei of the distal tip cell (arrow head), gonad sheath cells (arrows), and large intestinal nuclei (wedges). Several other gonadal sheath nuclei are not captured in the 8 μm-thick slice (see panel E). Scale bar: 100 μm.D. Negative control gonad preparation of the same TRA-2:HA strain as in C, but omitting the anti-HA primary antibody. Arrowhead marks the distal tip cell. Scale bar: 100 μm. Faint background in the germ line is comparable to that in A. E. Higher-magnification, 3 μm-thick slice of an HA-positive seam cell nucleus (arrowhead) and adjacent germ cell nuclei from the same gonad in A (alternative focal plane, near the bend, displayed rotated 90 degrees).

We next examined the TRA-2B:HA gonad expression in the *fog-2* null and *gld-1* null backgrounds. No clear elevation of TRA-2::HA signal was observed in the germ line of adult or L4 larval *fog-2(q71)* animals relative to wild type (**Fig. 6A**). Meiosis of XX spermatocytes begins in L3 (Hansen et al., 2004), it is likely sexual fate is also specified at this time. We therefore also compared wild-type and *fog-2(q71)* backgrounds at this stage. Anti-HA germline fluorescence was quantified by confocal microscopy and image analysis for 12 L3 gonads of each genotype. Again, no difference in expression was observed (mean pixel intensity of 41.17 vs. 41.2, two-tailed *t* Test, P=0.99). Thus, the repression of TRA-2 expression provided by *fog-2* is insufficient to raise the germ cell immunofluorescence signal above background levels when eliminated. Loss of *fog-2* does not alter the spatial expression of a GFP reporter for *oma-1* (Lin, 2003), a conserved direct GLD-1 target (Beadell et al., 2011; Lee and Schedl, 2004) easily detected in germ cells (**Fig. 6B**). We conclude that expression of mRNAs targeted by GLD-1 is not systematically up-regulated in the absence of FOG-2.

**Figure 6.**
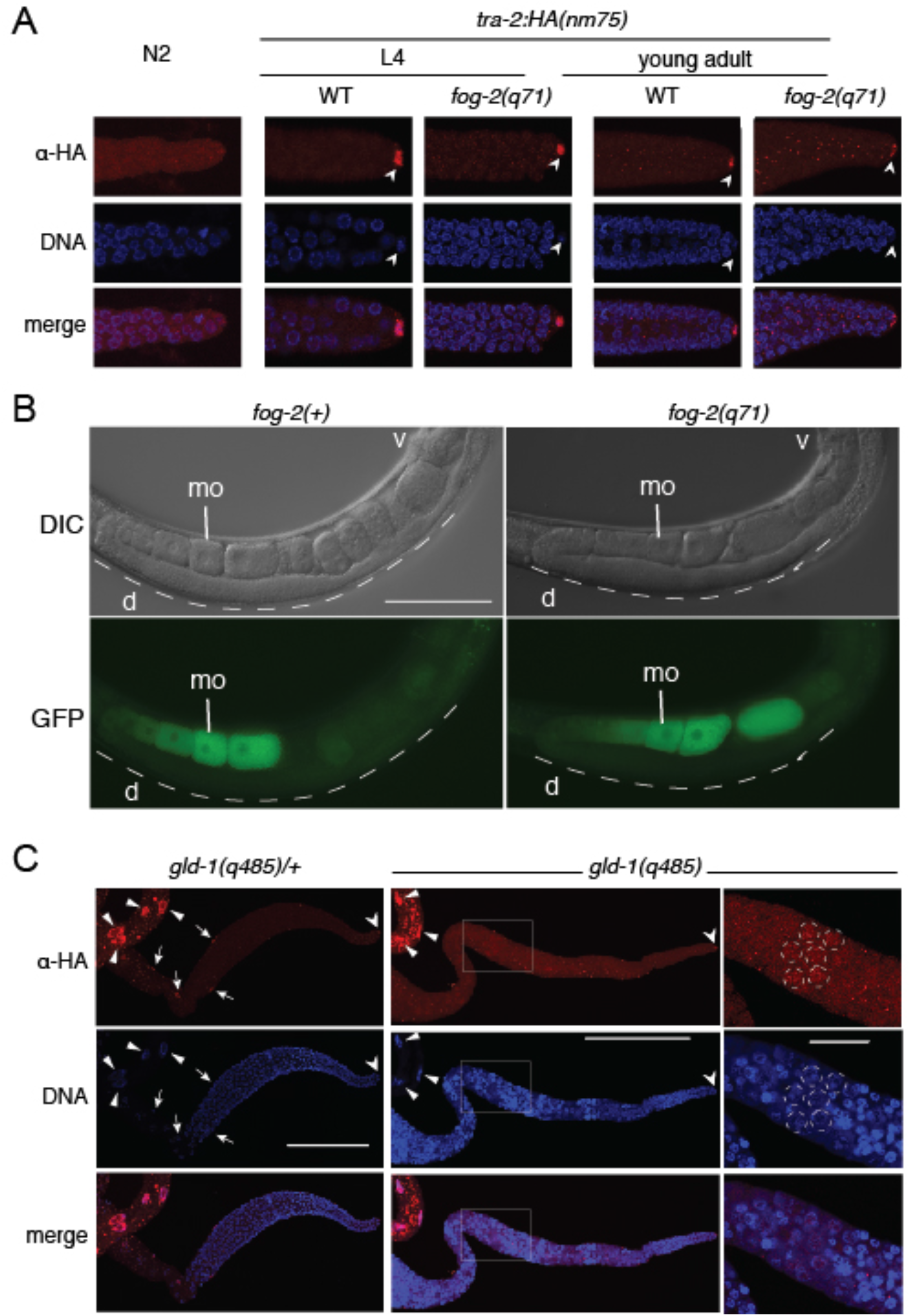
TRA-2 expression in *fog-2* and *gld-1* null mutants. A. Comparison of TRA-2B:HA expression in the distal region (typical of gonad as a whole) of otherwise wild-type *nm75* gonads (“WT”) and in *fog-2(q71)* mutants. Confocal micrographs of L4 larvae and young adults are shown for each, along with non-tagged N2 as a negative control (far left). The arrowheads indicate staining of the distal tip cell nucleus. Bright punctae in the *fog-2(q71)* adult sample (far right) were not consistently associated with a particular genotype from experiment to experiment. B. Differential interference contrast (top) and epifluorescence (bottom) micrographs of the *oma-1:GFP* reporter *teIs1* in live, anaesthetized wild-type (left) or *fog-2(q71)* homozygotes (right). GFP fluorescence is abundant in mature oocytes (mo), but lacking in both cases in the distal germline region (d, dashed line) where GLD-1 is expressed (Jones et al., 1996). Scale bar = 100 μm. C. Confocal micrographs of dissected gonads of *tra-2(nm75)* animals either heterozygous (left) or homozygous (right) for the *gld-1(q485)* null mutation, stained for TRA-2B:HA and imaged under identical conditions in the same session. Each image represents maximum intensity projections of 12-14 μm, or about half the depth of each gonad. As in Fig. 4 and panel 5A above, arrowheads, arrows, and wedges indicate the HA-positive distal tips cell, sheath cells, and gut nuclei, respectively; sheath cell nuclei in *gld-1(q485)* gonad are not visible in this focal plane. The far right column is an enlarged version of the region boxed in the tumorous *q485* gonad, a single focal plane with the red channel pixel intensity rescaled uniformly to highlight nuclear accumulation in a subset of pachytene nuclei (dashed circles). The scale bars in the left and middle columns are 100 μm, and 20 μm in the right column.

For *gld-1(q485)* mutants, immunofluorescence in the red (anti-HA) channel was elevated (roughly two-fold) compared to non-tumorous *gld-1(485/+)* siblings (**Fig. 6C**). This signal is mostly non-localized, but above this uniform background pachytene nuclei often show localized fluorescence. This suggests that there may be some up-regulation of TRA-2B in *gld-1(q485)* mutant germ cells. However, *gld-1(q485)* gonads are a qualitatively different tissue than wild-type or *q485/+* gonads, being composed of a vast excess of mitotically proliferating germ cells.

We therefore considered the possibility that their elevated fluorescence was due to intrinsically higher nonspecific background. To address this, we quantified the mean pixel intensity in sheath cell-free regions of *gld-1(q485)* gonads that were either processed normally, or without the anti-HA primary antibody to reveal nonspecific background. Processed and imaged in parallel on the same day, fluorescence of samples with anti-HA primary antibody (mean pixel intensity 12.69, S.E.M. 0.32, N=7) was significantly higher (unpaired two-tailed *t* test, P<0.0001) than of those that did not (mean pixel intensity 8.34, S.E.M. 0.47, N=4). The magnitude of the difference (1.5- fold) is similar to that seen between *gld-1(q485/+)* and *gld-1(q485)* samples stained with both primary and secondary antibodies. We conclude that loss of *gld-1*, unlike loss of *fog-2*, is sufficient to elevate TRA-2B expression to detectable levels in germ cells. The magnitude of the change is not large enough to be revealed in immunoblots of whole-worm lysates.

### *Sensitivity of tra-2(e2020) to loss of fog-2* and *gld-1 activity*

The *tra-2* gain-of-function *(gf)* alleles eliminate some or all GLD-1 binding sites in the 3’ UTR (Goodwin et al., 1993; Jan et al., 1999), and also eliminate XX spermatogenesis (Doniach, 1986). Such mutants can be restored to self-fertility by combination with a suitable *fem-3(gf)* allele, which on its own fully masculinizes the germ line (Barton et al., 1987). Schedl and Kimble (1988) found that in this context, seven different *tra-2(gf)* alleles tested were sensitive to *fog-2* loss. However, none of these alleles eliminate all known GLD-1 3’ UTR binding sites, suggesting continued sensitivity may be due to residual repression mediated by remaining sites. The *tra-2* gain-of-function *(gf)* allele *e2020* has a 108 nt deletion of its 3’ UTR that eliminates all GLD-1 binding *in vitro* (Goodwin et al., 1993; Jan et al., 1999). If FOG-2 regulates germline sex solely through its regulation of the *tra-2* mRNA via GLD-1 and its 3’ UTR binding sites, then *tra-2(e2020)* may be uniquely insensitive to *fog-2* loss. Conversely, feminization of the *e2020* allele by *fog-2(lf)* would suggest a role for *fog-2* that does not rely on the known *tra-2* 3’ UTR binding sites for GLD-1.

JK992 *(tra-2(e2020); fem-3(q95))* animals are normally Mog at 25° C, thus creating a situation in which abundant sperm (and no oocytes) are made without the known GLD-1-binding sites in the *tra-2* 3’ UTR. To assess whether *fog-2* has an impact on germline sex in this situation, we produced a *tra-2(e2020), fem-3(q95)* population that was also segregating for the *fog-2(q71)* null allele. Similar to results using weaker *tra-2(gf)* alleles with residual GLD-1-binding capacity (Schedl and Kimble, 1988), *tra-2(e2020gf); fem-3(q95gf); fog-2(q71)* triple mutants were all Fog (Table 2). However, their *fog-2(q71/+)* or *fog-2(+/+)* siblings remained Mog, with the exception of one heterozygote. Thus, lack of all *fog-2* function feminizes the germ line even when the *tra-2(gf)* allele used eliminates all known binding sites of GLD-1 in the *tra-2* 3’ UTR.

The above result could indicate that the FOG-2-GLD-1 complex has previously unrecognized targets, or that FOG-2 has GLD-1-independent functions. If loss of *gld-1* function in the same *tra-2(e2020gf); fem-3(q95gf)* context also feminizes, it would support the former explanation. We therefore performed an RNAi knock-down of *gld-1* in JK992 via injection (**Table 3**). This produces a highly penetrant oocyte tumor (Tum) phenotype in wild-type animals at both 15° and 25° C, and in JK992 at the permissive (oocyte-producing) 15° C. At 25°, *gld-1(RNAi)* shifted 12% of adults from Mog to Tum, indicating partial feminization. This is qualitatively similar to the effect of *fog-2(q71)*, but less penetrant. Whether this difference is due to incomplete knockdown or a GLD-1-independent function of FOG-2 remains unclear.

To more directly assess a possible role for FOG-2 in the absence of GLD-1, we sought to compare a sperm-producing *gld-1(lf)* to *gld-1(lf), fog-2(lf)*. The *fem-3(q95)* dominant Mog allele enables such a test, as *fem-3(q95); gld-1(q485)* double mutants are over 90% Mog at nonpermissive temperature (Francis et al., 1995b). We knocked down *gld-1* in *fem-3(q95); fog-2(q71)* double mutants (Table 3), and found that over half of the progeny of injected mothers were Tum. This indicates that loss of *fog-2* enhances the partial feminization of *fem-3(q95)* imparted by loss of *gld-1*.

### Dominant GLD-1 gain-of-function mutant exhibits weaker RNA-binding capacity

GLD-1 mutant alleles are categorized into 6 phenotypic classes (Francis et al., 1995a). The subclass C1 alleles, *q93, oz17, oz34, oz35*, and *q62* change Gly-248 into Arg (G248R); subclass C2 alleles, *oz30, oz16, oz29, oz33*, and *oz70* have a point mutation of G250R. The glycine residues altered by these missense mutations are highly conserved in the GSG domain (Jones and Schedl, 1995). XX animals from both sub-classes over-produce sperm and fail to switch to oogenesis (the masculinization of germline, or Mog phenotype). This has been interpreted as a gain-of-function phenotype, and alleles from both subclasses dominantly suppress the Fog phenotype of *fog-2(q71)* (Francis et al., 1995b).

One possible mechanism of a dominant *gain-of-function* GLD-1 mutant would be tighter binding to its target mRNA, which could suppress the translation to a greater degree. To test this possibility, we examined the GLD-1 wild type, G250R and G248R mutants for their binding affinity to a synthetic fragment encoding the 3’ UTR of *glp-1*, a GLD-1 target mRNA with a single binding site, using gel mobility shift assays. We found that while wild type GLD-1 bound to its target efficiently, the two mutant proteins had reduced binding (**Figure 7**). The G248R mutant had particularly poor binding. The C1 alleles bearing this mutation show a more severe Mog phenotype than the C2 alleles (G250R). Thus, contrary to one simple hypothesis, lower binding capacity of GLD-1 to its target mRNA is associated with a stronger dominant *gain-of-function* phenotype.

**Figure 7.**
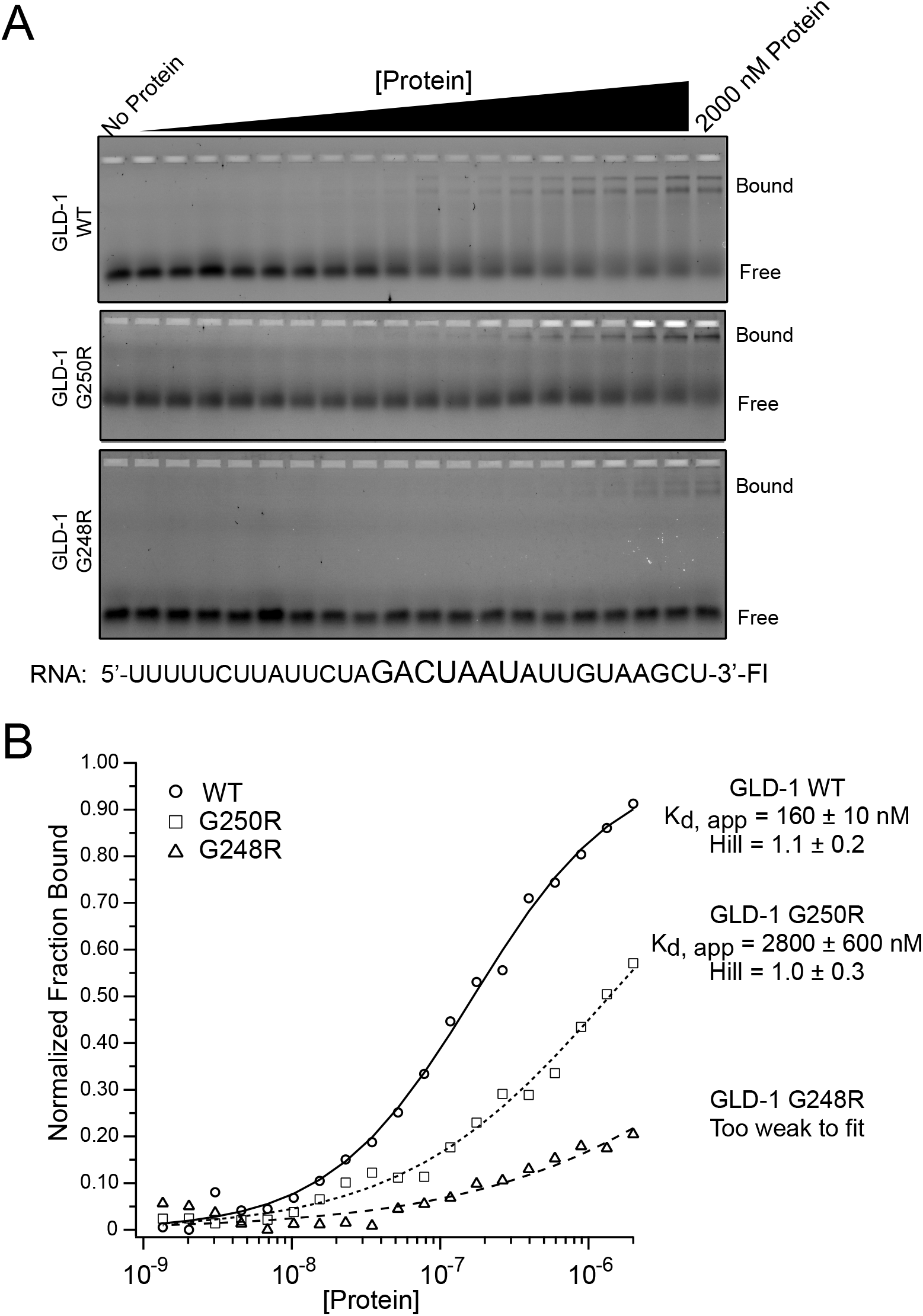
Binding analysis of GLD-1 WT, GLD-1 G248R, and GLD-1 G250R. A. An increasing concentration of purified, recombinant MBP-tagged GLD-1 or mutant variants was incubated with a fluorescein labeled RNA oligonucleotide comprising a fragment from the *glp-1* 3UTR that contains a GLD-1 binding motif, shown in larger font. Bound RNA was resolved from free using a slab polyacrylamide gel, and visualized using a FUJI FLA9000 Imaging System. B. The fraction of bound RNA was quantified using MultiGauge software. These data were plotted as a function of the protein concentration and fit to the Hill Equation in order to determine the apparent equilibrium dissociation constant (K_d_, _app_) and the Hill coefficient. The average and standard deviation of the K_d_, _app_ and the Hill equation for at least three experimental replicates are presented to the right of the graph. Data were normalized to the local minimum and the fitted window of GLD-1 WT binding to facilitate comparison of the three mutants on the same set of axes. Un-normalized data were used to obtain the fitted parameters. For GLD-1 G248R, the data were not fit because little to no binding was observed across three replicates.

## Discussion

### fog-2 and tra-2 regulation

The *fog-2* gene was formed in the *C. elegans* lineage by duplication and modification of one of many F-box proteins (Nayak et al., 2005). In contrast, *tra-2* and *gld-1* are ancient regulators of overall sex determination and oocyte development (Beadell et al., 2011; Haag and Kimble, 2000; Kuwabara, 1996). FOG-2 is thus a new player in an old system, and its recent origin and its essential role in the newly acquired self-fertility of *C. elegans* are likely not coincidental. By what mechanism does FOG-2 contribute to the regulation of germline sex in the *C. elegans* hermaphrodite? An important first step is to examine potential impacts on mRNA abundance. The transcriptome of *fog-2(q71)* XX gonads was previously characterized, and compared to those of *fem-3(q96)* mutants, which express only sperm (Ortiz et al., 2014). This comparison highlighted the extreme differences between the male and female germ cell transcriptomes, and revealed reductions in *tra-2* and *gld-1* expression in *q96* germ cells consistent with a negative feedback between sperm fate and their expression. However, to understand the role of FOG-2 in an otherwise female germ line, the comparison of *fog-2(q71)* with wild-type hermaphrodites is more appropriate.

Using both oligo-dT primed and random-primed libraries, we found that the expression of 99% of detected transcripts was unchanged in *fog-2* mutants relative to wild-type hermaphrodites. Similarly, nearly all GLD-1 target mRNAs, including *tra-2*, were also unchanged in their abundance. This finding is consistent with the hypothesis that FOG-2 regulates *tra-2* activity post-transcriptionally. It should be noted, however, that in our adult samples the XX spermatocytes that rely upon *fog-2* for their formation are not present. L4 samples would include such cells, but they would also likely include many spermatogenesis-related transcripts in the wild-type sample that would dominate the set of differentially expressed genes. The choice to analyze adults is somewhat justified because FOG-2 is expressed in all stages of development (Clifford et al., 2000).

Although neither the wild-type nor *fog-2(q71)* animals compared with RNAseq were undergoing active spermatogenesis, we nevertheless found that expression of 14 major sperm protein (msp) paralogs was reduced in *fog-2* mutants. One possible explanation is that cryptic misexpression of sperm genes occurs in oocytes, and this is reduced (feminized) in *fog-2(q71)*. Alternatively, because the *fog-2* mutant hermaphrodites were inseminated by *fog-2(q71)* males, the difference may reflect residual transcripts in spermatozoa. It would not be surprising if *fog-2* male sperm were cryptically feminized, and thus transcribed *msp* genes at lower levels. This could be tested in the future.

Assuming that FOG-2 does indeed regulate translation, the mechanism by which this would occur remains elusive. Many F-box proteins have roles in marking target proteins for ubiquitination and subsequent proteasomal degradation, and FOG-2 interacts with the *C. elegans* Skp1 homolog SKR1 (Nayak et al., 2005). What might the target of a putative FOG-2-mediated ubiquitination be? It is unlikely to be GLD-1, as both are required for the same process. One interesting possibility is that, by virtue of being tethered to the *tra-2* 3’ UTR via GLD-1, FOG-2 targets the nascent TRA-2 peptide itself. Such co-translational modification via a 3’ UTR scaffold has been reported for the human protein CD47 (Berkovits and Mayr, 2015).

### Tissue-and sex-specificity of TRA-2 expression

Three other groups have examined the subcellular, tissue-level, and sex-biased expression of TRA-2. Lum et al. (2000) detected a transgene fusion of TRA-2B to GFP in somatic nuclei. The multi-copy array used would not be expected to express in germ cells due to silencing. However, using a polyclonal antibody, endogenous TRA-2B was similarly reported in intestinal nuclei of XX hermaphrodites, but not detected in male intestine or germ cells from both sexes (Shimada 2006). Finally, an integrated 3xFlag-tagged TRA-2 transgene was expressed in nuclei of the head (Mapes 2010), and could be detected in males as long as it was heterozygous (reporter homozygotes were feminized). Our immunoblots and immunohistochemistry of HA-tagged endogenous TRA-2 are broadly consistent with these earlier studies. We observed expression of TRA-2B in XX hermaphrodites, but could not detect it in germ cells, nor in males. The full-length 170 kDa TRA-2A isoform did not appear on our blots. This could be because it does not readily solubilize in sample buffer, but the absence of plasma membrane expression in stained tissues is more consistent with rapid cleavage by the TRA-3 protease in XX animals. Weak germline expression of TRA-2B is surprising, given available *in situ* data (Kohara, 2005) and extensive evidence suggesting cell-autonomous function in germ cells. This includes a 1.8 kb transcript encoding TRA-2B that is abundant in oocytes (Kuwabara et al., 1998). It is consistent, however, with extensive translational repression of the *tra-2* mRNAs in germ cells, and this is well documented (Goodwin et al., 1993; Jan et al., 1999). The modest upregulation of TRA-2B signal in the germ line of *gld-1* mutants (**Fig. 6C**) indicates that complete loss of GLD-1 does have a measurable effect. In contrast, *fog-2* either exerts a more subtle influence on TRA-2 expression than does *gld-1*, or it regulates *tra-2* independently of its steady-state protein product abundance. The model of *tra-2* regulation developed here (**Table 1, Fig. 4**) suggests that a scenario in which *fog-2* exerts a weaker effect on *tra-2* activity than *gld-1* is consistent with current genetic data. However, a full assessment of the processes that regulate *tra-2* activity awaits more sensitive methods.

Though undetectable in wild-type germ cells via immunohistochemistry, TRA-2B:HA is highly expressed in distal tip cells and gonad sheath cells, somatic cells tightly directly attached to the germ line. This raises the possibility that the somatic TRA-2 expression may play a role in determining the sex of germ cells. Ablations of sheath cells can feminize *C. elegans* hermaphrodites (McCarter et al., 1997). Because sheath cell TRA-2 expression would be expected to promote oocyte fate, the possible connection between this and the sheath cells’ role in supporting XX spermatogenesis remains mysterious. Preliminary experiments in which the male-promoting protein FEM-3 was overexpressed in the distal tip cell of hermaphrodites led to abnormal somatic gonad morphology, but failed to masculinize germ cells (S.H. and E.S.H., unpublished data).

### Uncharacterized roles of fog-2 and gld-1 in germline sex determination

We present two surprising findings about the roles of *fog-2* and *gld-1* in the regulation of germline sex determination. First, the *tra-2(e2020); fem-3(q95); fog-2(q71)* and *tra-2(e2020); fem-3(q95); gld-1(RNAi)* triple mutants (**Tables 2 and 3**) indicate that loss of either *fog-2* or *gld-1* function can feminize germ cells in background that eliminates all known GLD-1-interacting motifs of the *tra-2* 3’UTR. This result reveals an unexpected aspect of their function. One possibility is that the 108 nucleotide deletion in the *tra-2(e2020)* 3’UTR does not abolish *in vivo* GLD-1 binding completely. For example, GLD-1, along with FOG-2, may bind the *tra-2* mRNA outside of the 3’ UTR. Some GLD-1 targets are repressed by binding in 5’ UTRs (Jungkamp et al., 2011), and a GLD-1 site in the *tra-2* open reading frame was predicted computationally (Brummer et al., 2013). Such an association may be enough to render sexual fate of *e2020* mutants sensitive to *fog-2* loss. Another possibility is that *fog-2* may be able to regulate TRA-2 expression in a GLD-1-independent manner, perhaps via its F-box domain. Finally, *fog-2* may promote sperm fate in an entirely TRA-2-independent manner, with or without interacting with GLD-1.

Second, the results in **Table 3** suggest that the sperm-promoting effects *gld-1* and *fog-2* are additive and at least partly independent. In the context of the *fem-3(q95)* dominant Mog mutation, *gld-1(lf)* suppresses masculinization less than 10% of the time (Francis et al., 1995b). However, *gld-1(RNAi); fem-3(q95); fog-2(q71)* animals are feminized (Tum) over half of the time, even though RNAi is unlikely to impart as a loss of *gld-1* function that is as strong as the *q485* mutation. That loss of *fog-2* enhances the Tum phenotype of *gld-1* indicates that FOG-2 has a sperm-promoting activity that is independent of GLD-1. This is unexpected, as recruitment of FOG-2 to the *tra-2* mRNA would seem to be dependent upon interaction with the GLD-1 RBP. Known FOG-2 protein interactors are GLD-1 (Jan et al., 1999) and the Skp1 homolog SKR-1 (Nayak et al., 2005), but these results suggest there may be others. Characterization of novel FOG-2-associated macromolecules is thus a priority for future research.

### Mechanism of translational repression by GLD-1

Transcriptome-wide examination of mRNA translation via ribosomal profiling found that many GLD-1 targets are translationally repressed (Scheckel et al., 2012). The feminized germline phenotypes of both *gld-1(lf)* and *tra-2(gf)* mutants are consistent with such negative regulation of TRA-2 expression by GLD-1. Our observation of enhanced immunofluorescence of TRA-2B:HA in *gld-1(q485)* gonads (**Fig. 6B**) provides further support for this. Dominant *gld-1(Mog)* alleles with the opposite phenotype as *gld-1* null mutants might therefore be interpreted as “hyper-repressors” of *tra-2*. However, we find that the missense mutant proteins these alleles encode actually bound the *tra-2* 3’ UTR more weakly *in vitro* than wild-type GLD-1.

How could poor binding lead to a phenotype that appears to represent a gain of function? Previous studies have suggested that GLD-1 not only represses translation, but also protects its targets from nonsense-mediate decay and perhaps other forms of degradation (Lee and Schedl, 2004; Scheckel et al., 2012). Thus, a mutation that weakens binding might have the counterintuitive effect of actually hyper-repressing *tra-2* expression, due to destabilization of the mRNA. Consistent with this, *tra-2* mRNA is slightly less abundant in *gld-1* mutants (Scheckel et al., 2012). However, other observations suggest the situation is not so simple. First, both *gld-1* null mutations and deletion of the SBE/GBMs of *tra-2* in the *e2020* allele have the opposite (Fog) phenotype of these *gld-1(gf)* alleles (Doniach, 1986; Francis et al., 1995b; Goodwin et al., 1993). Disruption of GLD-1-mRNA interaction via the C-class Mog alleles of GLD-1 is therefore not equivalent to a general loss of *gld-1* function. Second, the dominant Mog phenotype of *gld-1(q93)* is suppressed to self-fertility by one copy of the *tra-2(gf)* allele, *q122* (Francis et al., 1995b). This is consistent with the existence of a mode of *tra-2* repression for GLD-1 that does not require (and may be blocked by) wild-type levels of RNA-binding activity. Alternatively, the *gld-1(gf)* alleles tested here may confer novel protein-protein interactions that more than compensate for reduced RNA-binding capacity. Which, if either, of these scenarios is correct awaits further characterization of GLD-1’s many roles in germ cell gene regulation and development.

## Acknowledgements

We thank Tim Schedl for advice and for sharing strain JK992, Amy Beaven for assistance with confocal microscopy, and two reviewers for perceptive suggestions. This research was supported by NSF award IOS-1355119 to E.S.H. and an Ithaca College H&S STEM Summer Scholar award to L.E.R. Some strains were provided by the *Caenorhabditis* Genetics Center at the University of Minnesota, which is funded by NIH Office of Research Infrastructure Programs (P40 OD010440).

